# Enteric pathogens induce tissue tolerance and prevent neuronal loss from subsequent infections

**DOI:** 10.1101/2021.04.09.439221

**Authors:** Tomasz Ahrends, Begüm Aydin, Fanny Matheis, Cajsa Classon, Gláucia C. Furtado, Sérgio A. Lira, Daniel Mucida

**Affiliations:** Laboratory of Mucosal Immunology, The Rockefeller University, New York, NY, USA; Laboratory of Development and Homeostasis of the Nervous System, The Francis Crick Institute, London, UK; Precision Immunology Institute, Icahn School of Medicine at Mount Sinai, New York, NY, USA

## Abstract

The enteric nervous system (ENS) controls several intestinal functions including motility and nutrient handling, which can be disrupted by infection-induced neuropathies or neuronal cell death. We investigated possible tolerance mechanisms preventing neuronal loss and disruption in gut motility after pathogen exposure. We found that following enteric infections, muscularis macrophages (MMs) acquire a tissue-protective phenotype that prevents neuronal loss and dysmotility during subsequent challenge with unrelated pathogens. Bacteria-induced neuroprotection relied on activation of gut-projecting sympathetic neurons and signaling via β_2_-adrenergic receptors (β2AR) on MMs. In contrast, helminth-mediated neuroprotection was dependent on T cells and systemic production of interleukin (IL)-4 and -13 by eosinophils, which induced arginase-expressing MMs that prevented neuronal loss from an unrelated infection located in a different intestinal region. Collectively, these data suggest that distinct enteric pathogens trigger a state of disease- or tissue tolerance that preserves ENS number and functionality.

## Introduction

The gastrointestinal (GI) tract needs to simultaneously generate tolerance to harmless or beneficial dietary and microbial antigens and resistance to pathogen invasion. Additionally, disease tolerance to pathogen- and inflammation-induced tissue damage must operate in the intestine to maintain homeostasis (Medzhitov et al., 2012; Soares et al., 2014). This is particularly relevant for cells with reduced proliferative or regenerative capacity, such as intrinsic enteric-associated neurons (iEANs), which are abundantly present in the gut and serve to modulate intestinal motility and secretory function (Balemans et al., 2017; Furness et al., 2013; O’Leary et al., 2019).

Previous studies in mice and rats, supported by clinical observations, indicate enteric infections can lead to neuronal loss, long-term dysmotility and neuropathies (Holschneider et al., 2011; Matheis et al., 2020; Ohman and Simren, 2010; White et al., 2018). Yet, whether a state of tolerance can be induced after exposure to pathogens, preventing cumulative neuronal loss and functional changes, is still not known.

We investigated possible tolerance mechanisms preventing neuronal loss and disruption in gut motility after pathogen exposure. We found that following helminth or bacterial infection and via distinct pathways MMs acquire tissue-protective qualities and prevent neuronal loss during subsequent challenge with an unrelated pathogen. Helminth-induced neuroprotection was dependent on T cells and systemic production of interleukin (IL)-4 and -13 by eosinophils and maintained long-term through the modulation of hematopoietic progenitors in the bone marrow. Conversely, bacteria-induced neuroprotection relied on β_2_-adrenergic receptor signaling in MMs. Collectively, these data suggest that distinct enteric pathogens trigger a state of tolerance aimed to preserve the numbers and functionality of the ENS.

## Results

### *Y. pseudotuberculosis* infection induces iEAN protection during subsequent infection

To test whether primary infection can prevent neuronal loss after secondary challenge with an unrelated pathogen, we used an attenuated strain of *Salmonella* Typhimurium, *spiB*, which harbors a mutation in the type-III secretion system, impacting its intracellular replication (Tsolis et al., 1999). In contrast to wild-type *Salmonella, spiB* is not lethal and is cleared from wild-type C57BL/6 mice in 7-10 days, inducing a less severe form of Salmonellosis; nevertheless, *spiB* infection is sufficient to induce rapid iEAN loss and long-term dysmotility (Matheis et al., 2020). We thus first asked whether primary infection with an unrelated bacterium influences the tissue’s response to a subsequent exposure to *spiB*. We used *Yersinia pseudotuberculosis* (*Yp*), which like *Salmonella*, is a foodborne Gram-negative bacterium that primarily localizes to the ileum (Fonseca et al., 2015), causing a transient infection undetectable in the feces by 21 days postinfection (dpi) (Figure 1A). Primary infection with *Yp* led to significant iEAN loss, however not as pronounced as after primary *spiB* infection. Additionally, as suggested by our recent studies (Matheis et al., 2020), *spiB* exposure following an earlier *Yp* infection did not result in further iEAN loss (Figure 1B). This was not a result of an improved resistance to *spiB* as mice previously infected with *Yp* displayed similar pathogen load and clearance pattern to naïve mice (Figure 1C).

**Figure 1.**
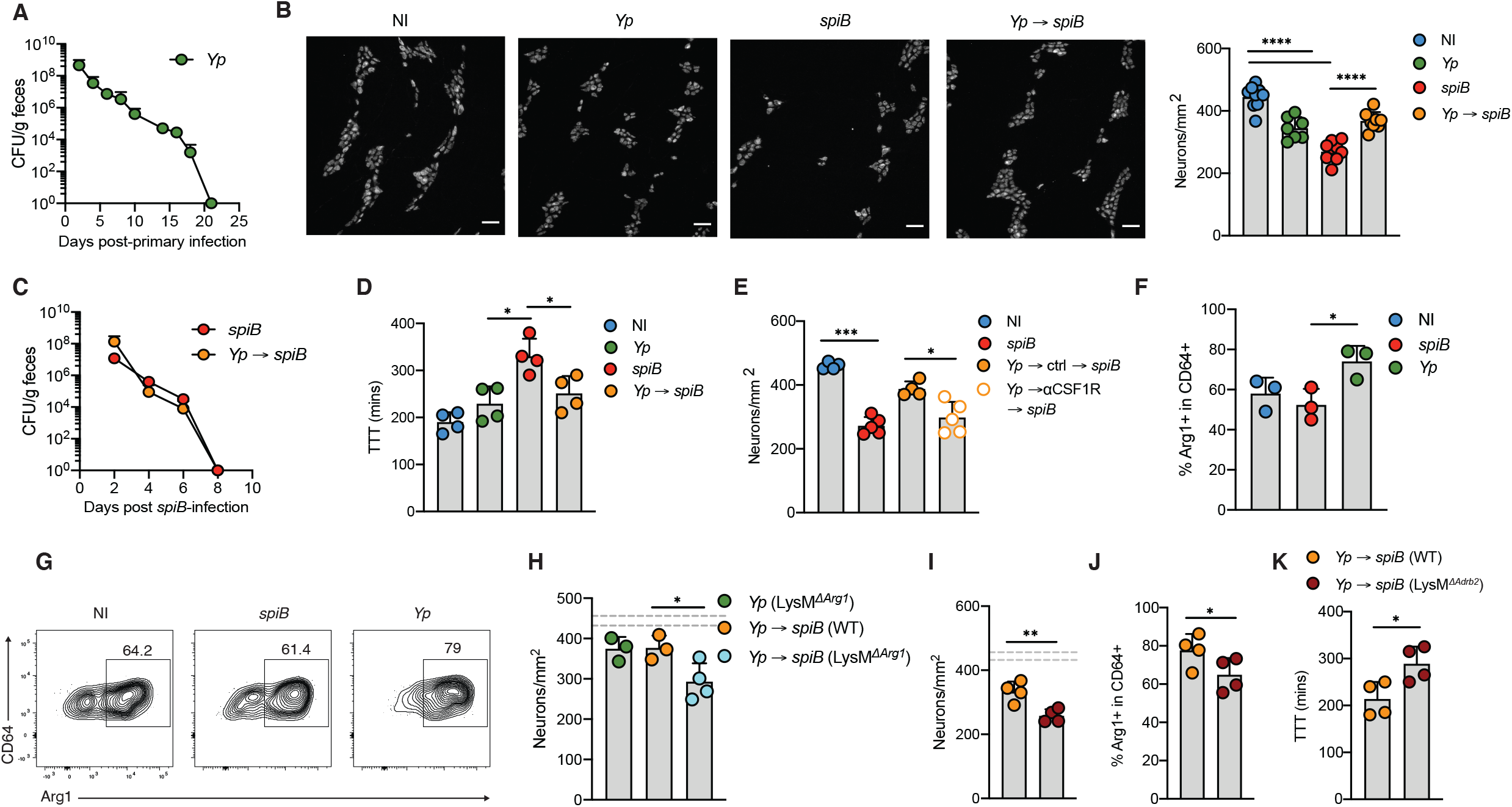
*Y. pseudotuberculosis* infection induces iEAN protection during subsequent infection. (A-D) C57BL/6J mice were orally gavaged with PBS (non-infected, NI), 10^9^ colony-forming units (CFU) of *Salmonella typhimurium spiB*, 10^8^ CFU of *Y.pseudotuberculosis* (*Yp*) only or 10^8^ CFU of *Yp* and 21 days later with 10^9^ CFU of *spiB*. (A) Quantification of fecal *Yp* CFU. (B) Neuronal quantification in the ileum myenteric plexus assessed by IF staining of ANNA-1 on day 10 post-*spiB* infection. Left, representative images; Scale bars, 50 μm. (C) Quantification of fecal *spiB* CFU. (D) Total GI transit time measured at 12 days post infection (dpi). (E) C57BL/6J mice were orally gavaged with PBS, *spiB* or *Yp*. At 20 dpi *Yp*-infected mice were treated with anti-CSF1R or IgG isotype control, followed by oral gavage with *spiB*. Ileum myenteric plexus neurons were quantified at 7 dpi. (F and G) Flow cytometry analysis of Arg1-expressing macrophages isolated from ileum muscularis at 14 dpi with *Yp* or *spiB*. (H) Quantification of ileum myenteric plexus neurons of LysM^*ΔArg1*^ mice and WT littermates orally gavaged with *Yp* only or followed by *spiB* infection 21 days later. (I-K) LysM^*ΔAdrb2*^ (Cre+) mice and WT (Cre-) littermates were orally gavaged with *Yp* and 21 days later with *spiB*. (I) Neuronal quantification of ileum myenteric plexus 7 days post-*spiB* infection. (J) Arg1 expression by macrophages in ileum muscularis at 7 dpi as determined by flow cytometry. (K) Total gastrointestinal (GI) transit time at 12 dpi. Dashed lines indicate the range of day 7 iEAN numbers defined by mean ± SEM of a large set of control C57BL/6J mice. Data is from 2 pooled independent experiments (3-5 mice per condition, A to C) or one representative experiment (D-K). Error bars indicate SD, * p ≤ 0.05, *** p ≤ 0.001, **** p ≤ 0.0001 (unpaired Student’s t test or ANOVA with Tukey’s post-hoc test).

Disease tolerance strategies alleviate the fitness costs by promoting resilience in the presence of an insult (Ayres, 2020). Because one of the main roles of the ENS in regard to fitness cost is the control of intestinal motility, we measured GI transit time as a functional readout for neuronal loss. Decreased iEAN numbers generally correlated with increased GI transit time; while *spiB* infection led to increased GI transit time, previous *Yp* infection preserved gut motility after subsequent *spiB* challenge (Figure 1D). These results point to development of tissue-, or disease tolerance (Ayres, 2020; Martins et al., 2019) post-bacterial infection, which prevented neuronal loss to subsequent infection with a different pathogen.

We have previously shown that MMs quickly acquire a tissue-protective phenotype and limit iEAN loss during primary *spiB* infection, a process that depends on β_2_ adrenergic receptor (β_2_-AR) signaling on MMs (Gabanyi et al., 2016; Matheis et al., 2020). Our studies also pointed to Arginase 1-dependent polyamine production by MMs, which is induced upon infection in a β_2_-AR-dependent manner, mediating a basal neuroprotective effect during primary infections (Gabanyi et al., 2016; Matheis et al., 2020). To test whether *Yp*-induced tolerance relied on MMs, we used anti-CSF1R antibody, which selectively depletes MMs while preserving lamina propria macrophages (LpM) when administered at 50 mg/kg (Matheis et al., 2020; Muller et al., 2014). Administration of anti-CSF1R antibody to mice post *Yp* clearance led to a pronounced iEAN loss following challenge with *spiB*, suggesting the requirement for MMs to maintain *Yp*-induced neuronal protection (Figure 1E). Flow cytometric analysis of the ileum from mice after primary infection revealed that *Yp* infection leads to the upregulation of Arg1 by MMs beyond a basal level (Figures 1F and 1G). To examine whether heightened Arg1 expression by MMs is required for *Yp*-induced neuroprotection, we used LysM^Δ*Arg1*^ mice, in which *Arg1* is conditionally deleted in myeloid cells. *Yp*-mediated neuroprotection was abolished in LysM^Δ*Arg1*^ following challenge with *spiB* infection (Figure 1H). Moreover, this neuroprotective mechanism also required b2-AR expression by MMs, as LysM^Δ*Adrb*2^ mice previously infected with *Yp* showed significant neuronal loss compared to wild-type littermate control mice upon *spiB* infection (Figure 1I). This effect correlated with decreased Arg1 expression by MMs and increased GI transit-time (Figure 1J and 1K). Consistently, while disease tolerance was impaired by both Arg1 and β_2_-AR targeting on MMs, *spiB* load was not affected by these strategies (Figures S1A and S1B). Taken together, these results indicate that primary bacterial infection results in a state of disease tolerance that mediates neuroprotection to subsequent infections, preserving intestinal motility.

### Helminth infections induce long-term iEAN protection during subsequent infection

Next, we asked whether infection-induced neuroprotection could be induced by other pathogens. In particular, we aimed to determine whether helminths, which co-evolved with mammals and typically induce a very distinct immune response to bacterial pathogens, could prevent neuronal loss. We chose to infect mice with *Strongyloides venezulenesis* (*Sv*), a parasitic nematode that causes an acute infection and primarily localizes to the duodenum (Esterhazy et al., 2019; Silveira et al., 2002). Initial *Sv* infection was cleared by 12 dpi and, contrary to the bacterial pathogens we tested, did not lead to iEAN loss in the ileum or duodenum (Figures 2A, 2B, S2A, and (Matheis et al., 2020)). However, primary *Sv* duodenum infection completely prevented iEAN loss in the ileum following subsequent *spiB* challenge (Figures 2B and S2A). This distal protection by previous, proximal *Sv* infection was not associated with changes in *spiB* load and resulted in the maintenance of normal gut motility (Figures 2C and 2D). Additionally, while the proximal intestine-helminth *Heligmosomoides polygyrus* (*Hp*) was recently shown to impair anti-neurotropic virus immunity (Desai et al., 2021), distal protection from iEAN loss in the ileum was also observed in *Hp*-infected and ivermectin-treated mice subsequently infected with *spiB* (Figure S2B). Moreover, initial *Sv* infection also led to neuroprotection after subsequent *Yp* challenge (Figure S2C). Consistent with a disease tolerance phenotype, primary *Sv* or *Hp* infection did not affect subsequent clearance of *spiB* or *Yp*, respectively (Figures S2D and S2E).

**Figure 2.**
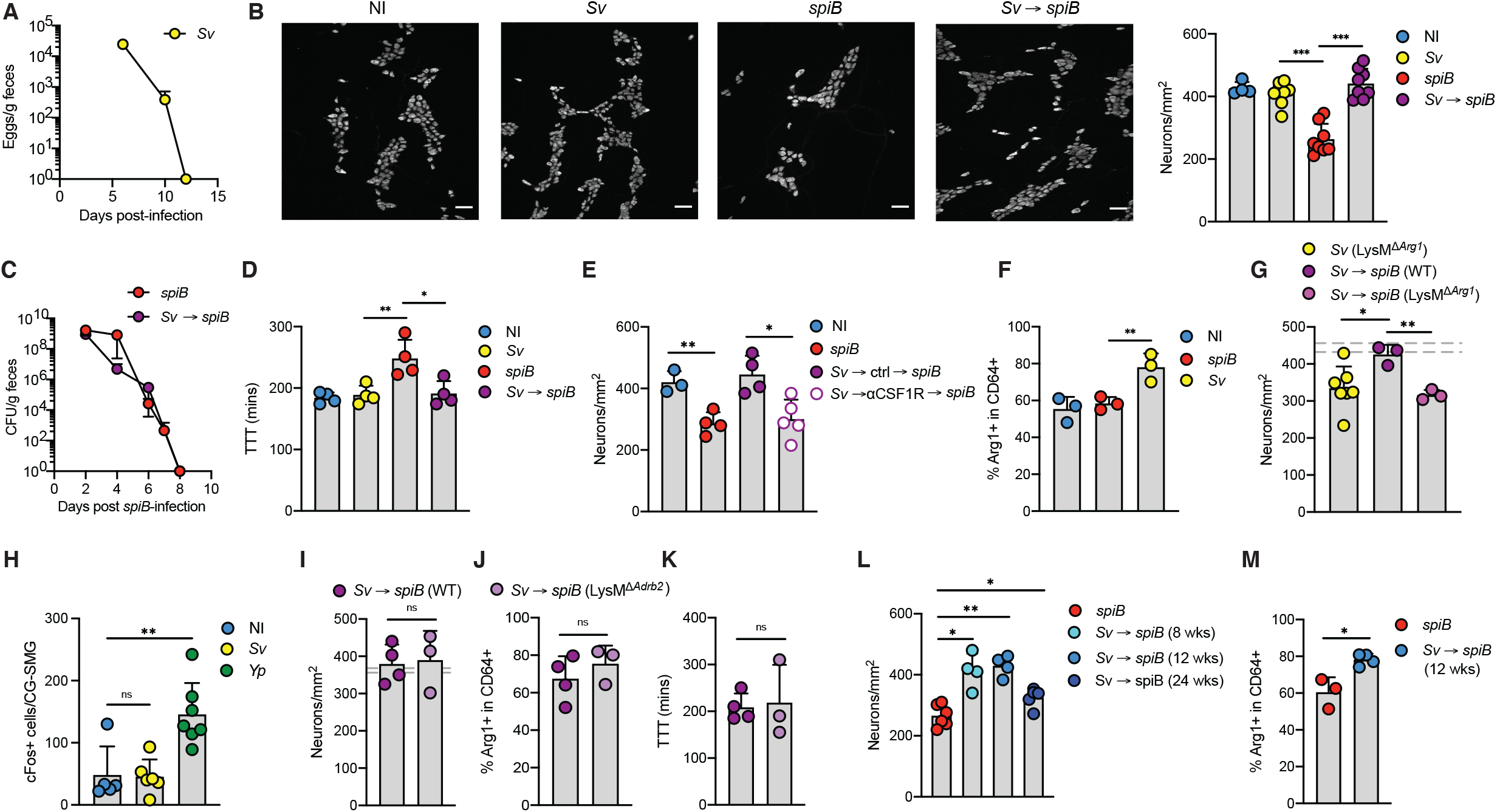
*S. venezuelensis* infection induces long-term iEAN protection during subsequent infection. (A-D) C57BL/6J mice were s.c. injected with water and then orally gavaged with PBS (non-infected, NI) or infected with 10^9^ colony-forming units (CFU) of *Salmonella Typhimurium spiB*, 700 *S. venezuelensis* (*Sv*) larvae only or 700 *Sv* larvae and 14 days later with 10^9^ CFU of *spiB*. (A) Quantification of fecal *Sv* eggs. (B) Right, Neuronal quantification in the ileum myenteric plexus assessed by IF staining using ANNA-1 on day 8 post-*spiB* infection. Left, representative images; Scale bars, 50 μm. (C) Quantification of fecal *spiB* CFU. (D) Total GI transit time measured at 10 dpi. (E) C57BL/6J mice were orally gavaged with PBS or infected with *spiB* or Sv. At 14 dpi *Sv*-infected mice were treated with anti-CSF1R or IgG isotype control, followed by oral gavage with *spiB*. Ileum myenteric plexus neurons were quantified 7 days post-*spiB* infection. (F) Flow cytometry analysis of Arg1-expressing macrophages isolated from ileum muscularis at 14 dpi with *Sv* or *spiB*. (G) Neuronal quantification of ileum myenteric plexus of LysM^*ΔArg1*^ (Cre+) mice and WT (Cre-) littermates orally gavaged with *Sv* only or followed by *spiB* infection 14 days later. (H) Number of cFos+ neurons in CG-SMG 3 days after *Yp* infection, 7 days after *Sv* infection and in non-infected controls. (I-K) LysM^*ΔAdrb2*^ mice and WT littermates were orally gavaged with *Sv* and 14 days later with *spiB*. (I) Quantification of ileum myenteric plexus neurons 7 days post-*spiB* infection. (J) Arg1 expression by macrophages in ileum myenteric plexus at 7 dpi. (K) Total GI transit time at 10 dpi. (L and M) C57BL/6J mice were orally gavaged with *spiB* only or *Sv* and 8, 12 or 24 weeks later with *spiB*. (L) Ileum myenteric plexus neurons were quantified 10 days post-*spiB* infection. (M) Arg1 expression by macrophages in ileum myenteric plexus at 7 dpi as determined by flow cytometry. (G and I) Dashed lines indicate the range of day 7 iEAN numbers defined by mean ± SEM of all control C57BL/6J mice. Data is from 2 pooled independent experiments (3-5 mice per condition, A-C and H) or one representative experiment (D-G and I-M). Error bars indicate SD, ns – not significant, * p ≤ 0.05, ** p ≤ 0.01, *** p ≤ 0.001 (unpaired Student’s t test or ANOVA with Tukey’s post-hoc test).

*Sv*-induced tolerance depended on MMs, as evidenced by anti-CSF1R antibody-mediated depletion of MMs post *Sv* clearance, which abolished the neuroprotection upon *spiB* infection (Figure 2E). In line with our observations post primary *Yp* infection, following *Sv* challenge ~80% of MMs expressed Arg1, compared to ~60% observed after *spiB* infection (Figures 2F and S2F). Of note, MMs in the duodenum expressed heightened Arg1 already at steady state compared to the ileum, and this was not further increased following *spiB* or *Sv* infection (Figure S2F). LysM^Δ*Arg1*^ mice previously infected with *Sv* showed significant neuronal loss following secondary *spiB* infection, in contrast to wild-type controls (Figure 2G). In opposition to *Yp*, *Sv* infection did not result in activation of gut-projecting sympathetic neurons (Figure 2H). Conversely, in contrast to the phenotype observed post *Yp*, *Sv*-mediated iEAN protection to subsequent *spiB* infection was maintained in LysM^Δ*Adrb2*^ mice. Mice lacking β_2_-AR signaling in myeloid cells previously infected with *Sv* also sustained similar iEAN numbers, Arg1 expression by MMs, and GI transit time upon *spiB* infection as wild-type littermate controls (Figures 2I-2K). Moreover, initial infection with *Sv* did not affect *spiB* load and clearance in either LysM^Δ*Adrb2*^ and LysM^Δ*Arg1*^ mice (Figures S2G and S2H). These results indicate that while converging on neuroprotective MMs, *Yp*- and *Sv*-induced disease tolerance mechanisms are distinct.

We also examined how long *Sv*-mediated neuroprotection is maintained after the initial infection is cleared. Subsequent *spiB* challenge up to 12 weeks after primary *Sv* infection did not result in any iEAN loss. Challenge at 24 weeks post-infection did result in significant iEAN loss, however the enteric neuron numbers were still higher when compared to controls infected with *spiB* only. (Figure 2L). Long-term iEAN protection correlated with sustained high expression of Arg1 in MMs observed up to 12 weeks post-infection (Figure 2M). These results reveal a long-term neuroprotection in the ileum induced by a single duodenal helminth infection.

### Immune response to *S. venezuelensis* is associated with iEAN protection during subsequent infection

To identify a mechanism responsible for the long-term helminth–driven disease tolerance, we next further characterized the immune response to *Sv*. Helminths are known to drive a robust type-2 immunity, with accumulation of innate immune cells, including tuft cells, eosinophils and group 2 innate lymphoid cells (ILC2), and later CD4^+^ T helper 2 (Th2) cells (Maizels, 2020; O’Leary et al., 2019; Vivier et al., 2018). Coinfections with helminths may result in deviation of appropriate immune responses resulting in breakdown of barrier and increased susceptibility to unrelated pathogens (Desai et al., 2021; Marple et al., 2017). We observed an accumulation of Gata3^+^ CD4^+^ “Th2” cells in the duodenum, but not in the ileum LP following *Sv* infection, which was further increased when mice underwent subsequent *spiB* infection (Figures 3A and S3A). Additionally, we found a significant increase in the frequencies of mucosal mast cells and eosinophils in the LP and intraepithelial (IE) compartments of the ileum and duodenum of *Sv*-infected mice (Figures 3B, 3C, S3B, and S3C). At steady-state, however, the duodenum harbored higher numbers of eosinophils than the ileum, the former reaching almost 80% of total CD45^+^ cells found in the LP post *Sv*-infection (Figures 3B and 3C). Of note, duodenal eosinophilia was maintained for up to 12 weeks post-infection and diminished at week 24 (Figure 3D), reflecting changes observed in MMs, GI transit time and neuroprotection post-*Sv* infection described above. Consistent with an overall heightened type-2 immunity, serum of *Sv*-infected mice showed prolonged increase in interleukin (IL)-4 and IL-13 levels (Figures 3E and 3F).

**Figure 3.**
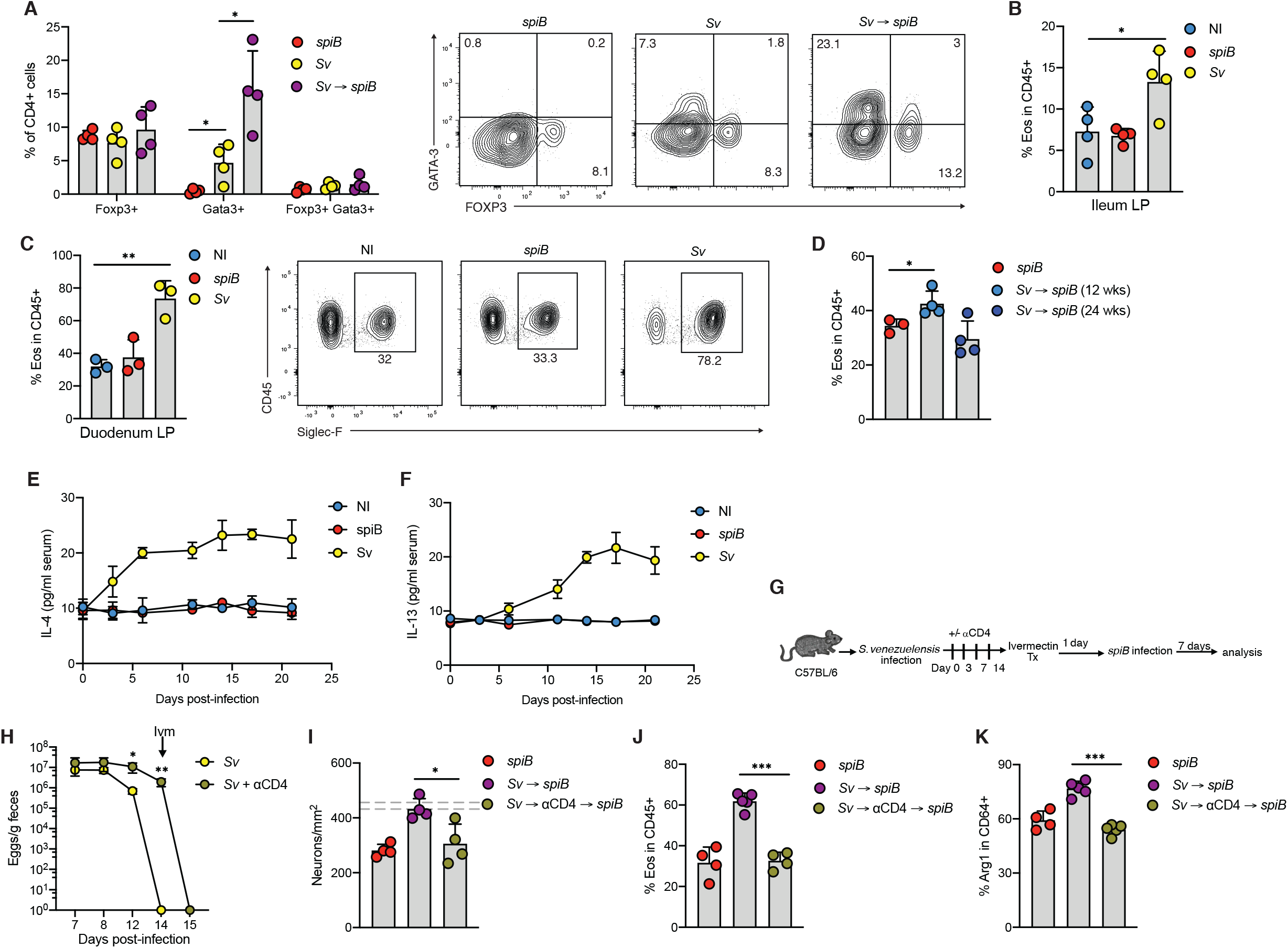
Immune response to *S. venezuelensis* is associated with iEAN protection during subsequent infection. (A-F) C57BL/6J mice were s.c. injected with water (non-infected, NI) or infected with *spiB, Sv* only or *Sv* and 14 days later with *spiB*. (A) Flow cytometric intranuclear analysis of CD4^+^ T cells in lamina propria (LP) expressing indicated transcription factors 2 days post-*spiB* infection. (B and C) Frequency of ileum (B) and duodenum (C) LP eosinophils at 14 dpi with *Sv*. (D) Frequency of duodenum LP eosinophils in mice infected with *spiB* only and *Sv* followed by *spiB* 12 or 24 weeks later and analyzed at 7 dpi. (E and F) Serum levels of IL-4 (E) and IL-13 (F) measured over time post-infection with *spiB* or *Sv*. (G) Experimental design for (H-K). (H-K) C57BL/6J mice were infected with *spiB* or *Sv*. On indicated days *Sv*-infected mice were treated with anti-CD4 antibody, followed by treatment with ivermectin and oral gavage with *spiB*. (H) Quantification of fecal *Sv* eggs. (I) Quantification of ileum myenteric plexus neurons 7 days post-*spiB* infection. Dashed lines indicate the range of day 7 iEAN numbers defined by mean ± SEM of a large set of control C57BL/6J mice. (J) Frequency of duodenum LP eosinophils at 7 dpi. (K) Arg1 expression by ileum myenteric plexus macrophages at 7 dpi. All data is representative of 2 independent experiments with 3-4 mice per condition. Error bars indicate SD, * p ≤ 0.05, ** p ≤ 0.01, *** p ≤ 0.001 (ANOVA with Tukey’s post-hoc test).

We next examined whether CD4^+^ T cells were required for *Sv*-induced neuroprotection using a depleting anti-CD4 antibody following primary infection (Figure 3G). Depletion of CD4-expressing cells hindered type-2 immunity and the capacity to clear *Sv* infection. Therefore, we treated mice with the anti-helminthic drug, ivermectin, before infecting them with *spiB* (Figure 3H). Mice treated with anti-CD4 antibody showed significant iEAN loss, which correlated with reduced eosinophilia and Arg1 expression by MMs (Figures 3I-3K). Tissue-resident memory T (T_RM_) cells mediate important *in situ* resistance functions and were also shown to provide cross-protection against different pathogens (Ariotti et al., 2014). We examined their role in *Sv*-induced neuroprotection by using mice with selective deficiency in the transcription factors Hobit and Blimp-1 (DKO mice), which were shown to be unable to generate T_RM_ cells (Mackay et al., 2016). Compared to wildtype controls, DKO mice displayed no differences in iEAN numbers, frequencies of eosinophils and Arg1-expressing MMs after successive exposure to *Sv* and *spiB*, suggesting that T_RM_ cells do not play a significant role in this process (Figures S3D-S3F). These results indicate that CD4^+^ T cells are required for *Sv*-induced neuroprotection, and also point to their “accessory” or “helper” role in this process.

We then assessed whether the two main innate immune cell types that accumulate during helminth infections, mast cells and eosinophils, play a role in *Sv*-induced neuroprotection. Mast cells accumulation after *Sv* infection was predominantly observed in the duodenum epithelial compartment (Figure S3B). To target mast cells, we generated a novel CRISPR-based *knock-in* strain in which human diphtheria toxin receptor (hDTR) and the td tomato fluorescent protein are expressed under a promoter preferentially active in mast cells (Lilla et al., 2011), carboxypeptidase A3 (*cpa3*). Administration of diphtheria toxin (DT) after initial *Sv* infection led to a significant reduction in mast cell numbers (Figure S3G). Consistent with a role for mast cells in resistance to helminths, DT-treated mice showed a significant delay in Sv clearance (Figure S3H). Nevertheless*, Sv*-infected DT-treated *cpa3*^DTR-tdTomato^ and control mice showed similar iEAN numbers upon subsequent *spiB* challenge, suggesting that mast cells are dispensable for neuroprotection (Figure S3I). To address a possible role for eosinophils in this process, we first treated *Sv*-infected mice with neutralizing anti-IL-5 antibody followed by *spiB* infection. ILC2- and Th2-derived IL-5 drives eosinophil activation and accumulation (Maizels, 2020; O’Leary et al., 2019; Vivier et al., 2018); consistently, IL-5 blockade resulted in impaired eosinophilia (Figure S3J). Additionally, *Sv*-infected mice treated with anti-IL-5 displayed a significant iEAN loss and reduced Arg1 expression by MMs upon subsequent *spiB* exposure (Figures S3K and S3L). This data points to an important role for eosinophils in neuroprotection and disease tolerance post helminth infection.

### Pet store mice are resistant to neuronal loss upon *spiB* infection

We next turned to potential implications of infection-induced neuroprotection in free-living or nonlaboratory organisms, which are exposed to more pathogens and sustain a higher immune activation than laboratory animals (Abolins et al., 2017). We asked whether increased exposure to pathogens in mice maintained in a pet store resulted in altered iEAN numbers at steady state and loss upon bacterial infection. At steady state, pet store mice displayed lower iEAN numbers per mm^2^ when compared to naïve specific-pathogen-free (SPF) C57BL/6 mice (Figure 4A), resembling what we observed in SPF mice previously infected with *Yp*. However, we observed that pet store mice had significantly longer small intestines (Figure 4B). To account for these morphological differences, we analyzed the numbers of neurons found in each ganglion. Pet store mice had similar numbers of neurons per ganglion when compared to non-infected SPF controls (Figure 4C). Nevertheless, infection with *spiB* resulted in a decrease of both total number and neurons per ganglia in SPF, but not in pet store mice, although the latter also cleared *spiB* two days earlier than average SPF mice (Figures 4A, 4C and 4D). We noted a wide range in the frequency of eosinophils in the LP of pet store mice, probably reflecting their varied infection history (Figure 4E). Additionally, pet store mice showed overall elevated levels of serum IL-4 and IL-13, and Arg1 expression by MMs (Figures 4F and 4G). Finally, consistently with the increased frequency of eosinophils in the tissue, analysis of bone marrow of pet store mice revealed that both hematopoietic progenitors (Lineage^-^Sca1^+^c-Kit^+^, LSK) cells and eosinophil progenitor (EoP) cells were enriched when compared to SPF controls (Figures 4H-4J). These results suggest that non-SPF mice, likely due to their infection history, acquire long-term protective mechanisms that regulate or prevent neuronal loss during infections occurring later in life.

**Figure 4.**
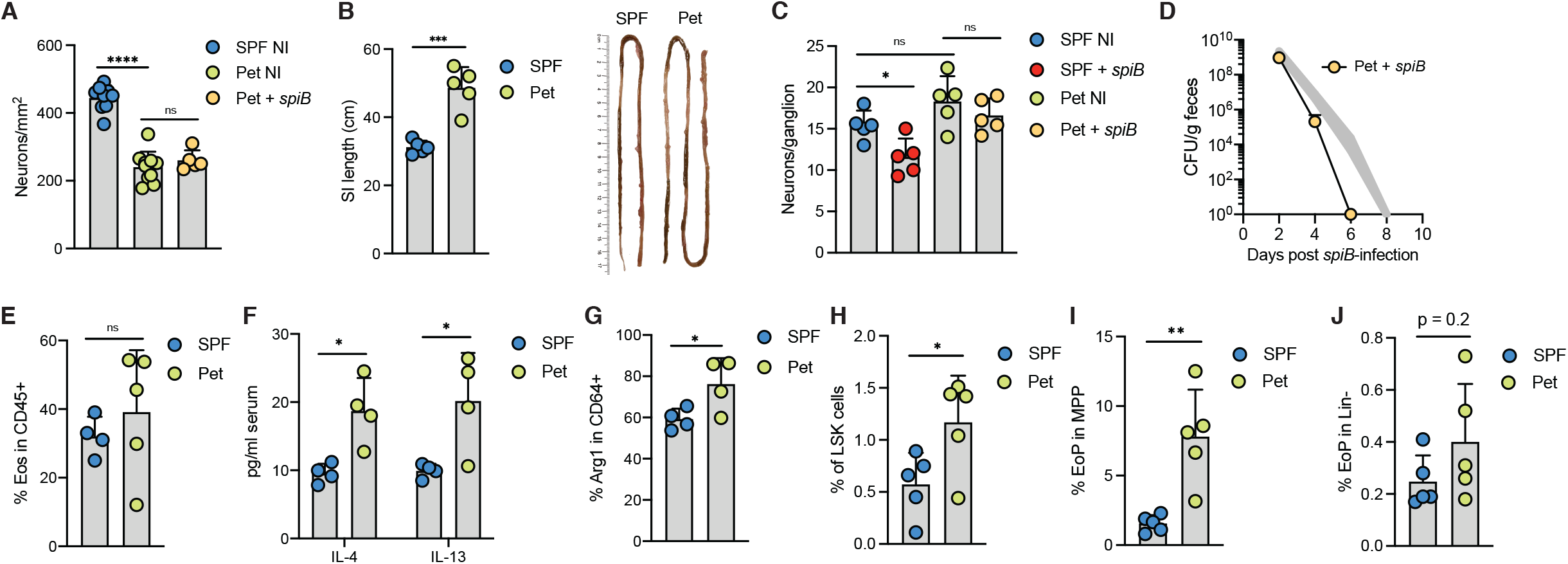
Pet store mice are resistant to neuronal loss upon *spiB* infection. Pet store mice were analyzed at steady-state (A-C and E-J) or 7 days post *spiB*-infection (A, C and D). (A) Quantification of ileum myenteric plexus neurons. (B) Small intestine length. (C) Number of iEAN per ganglion within the ileum myenteric plexus. (D) Quantification of fecal *spiB* CFU. Shaded area indicates range of *spiB* CFU numbers defined by mean ± SEM from a large set of control SPF mice. (E) Frequency of duodenum LP eosinophils. (F) Serum levels of IL-4 and IL-13. (G) Frequency of Arg1-expressing ileum myenteric plexus macrophages. (H) Frequencies of indicated bone marrow progenitor cell subsets. Data is from one representative (B-J) or 2 pooled independent experiments with 3-5 mice per condition (A). Error bars indicate SD, ns – not significant, * p ≤ 0.05, ** p ≤ 0.01, *** p ≤ 0.001, **** p ≤ 0.0001 (unpaired Student’s t test or ANOVA with Tukey’s post-hoc test).

### IL-4– and IL-13–producing eosinophils mediate *Sv*-induced neuronal protection

To directly investigate the contribution of eosinophils to this process, we utilized a genetic mouse model in which hDTR is expressed through eosinophil peroxidase locus (iPHIL) (Jacobsen et al., 2014). Administration of DT to *Sv*-infected iPHIL mice significantly reduced the numbers of eosinophils in duodenum and ileum LP (Figures 5A and 5B). Depletion of eosinophils did not affect egg load and *Sv* clearance, but abolished *Sv*-induced neuroprotection upon *spiB* infection (Figures 5C and 5D). This effect correlated with decreased expression of Arg1 by MMs and reduced GI motility (Figures 5E and 5F). In contrast, *Yp*-induced neuronal protection was not affected by depletion of eosinophils (Figures S4A and S4B).

**Figure 5.**
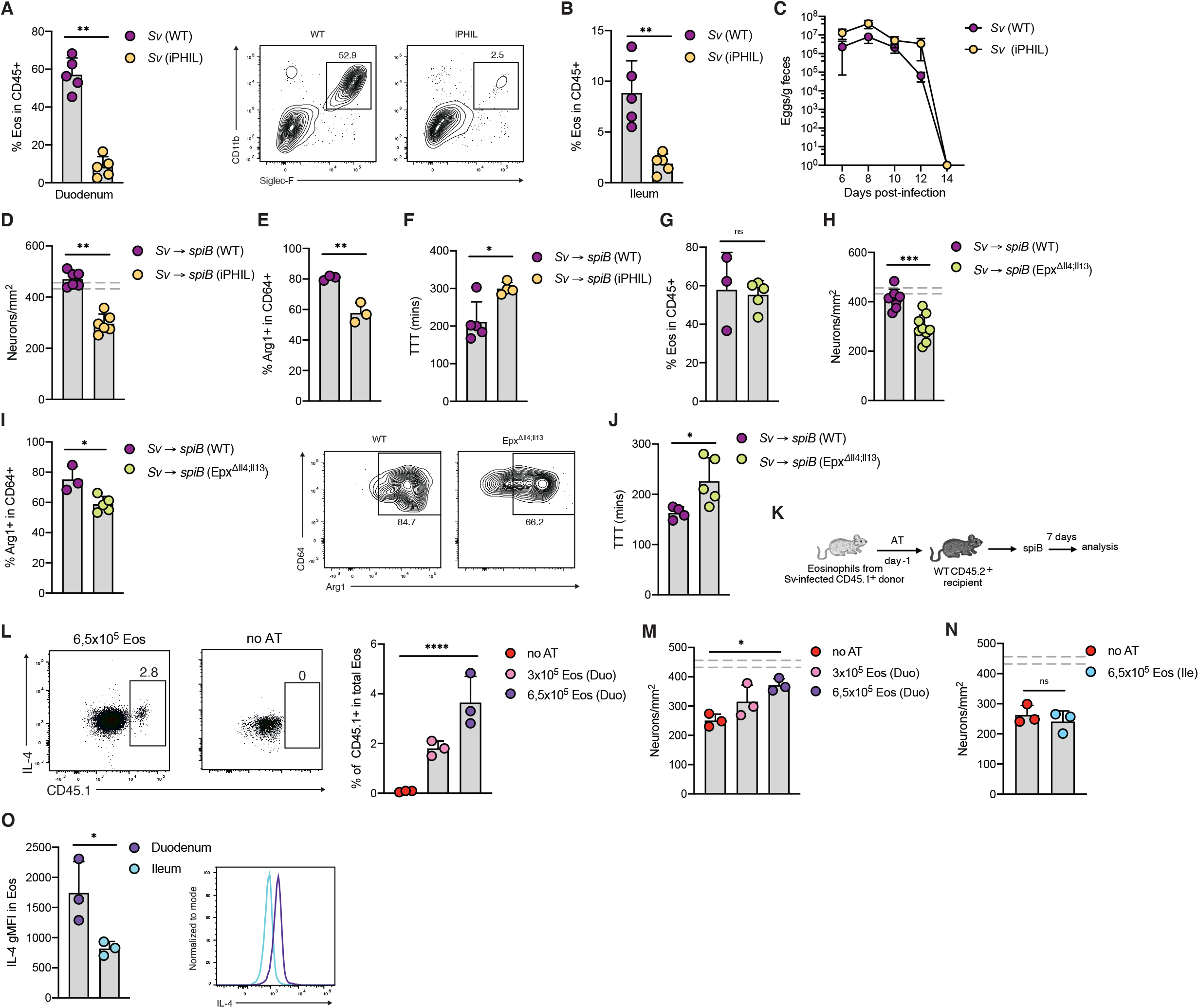
IL-4 and IL-13 producing eosinophils mediate *Sv*-induced neuronal protection. (A-C) iPHIL mice and WT littermates were infected with *Sv* and treated with DT on day 0, 1, 4, 6 and 8 post-infection. Analysis was performed at 10 dpi. (A and B) Frequency of duodenum (A) and ileum (B) LP eosinophils. (C) Quantification of fecal *Sv* eggs. (D-F) iPHIL mice and WT littermates were infected with *Sv* and treated with DT on day 0, 1, 4, 7, 10, 13, 16 and 18 post-infection. On day 14 post-*Sv* infection mice were orally gavaged with *spiB*. (D) Quantification of ileum myenteric plexus neurons at 7 dpi. (E) Arg1 expression by ileum myenteric plexus macrophages at 7 dpi. (F) Total GI transit time measured 10 days post-*spiB* infection. (G-J) Epx^Δ<u>Il4</u>;Il13^ mice or WT littermates were infected with *Sv* larvae and 14 days later with *spiB*. (G) Frequency of duodenum LP eosinophils at 7 dpi. (H) Quantification of ileum myenteric plexus neurons at 7 dpi. (I) Arg1 expression by ileum myenteric plexus macrophages at 7 dpi. (J) Total GI transit time measured 10 days post-*spiB* infection. (K) Experimental design for (L-N). (L-N) C57BL/6J CD45.2 mice received indicated numbers of eosinophils isolated from duodenum or ileum LP of CD45.1 mice infected with *Sv* 7 days earlier. Recipient mice were subsequently infected with *spiB*. (L) Frequencies of donor CD45.1^+^ duodenum eosinophils found in LP of recipient mice at 7 dpi. (M) Quantification of ileum myenteric plexus neurons in mice receiving duodenum (M) or ileum (N) eosinophils 7 days post-*spiB* infection. (O) C57BL/6J mice were infected with *Sv*. Expression of IL-4 was measured in duodenum and ileum eosinophils 7 at dpi. Dashed lines indicate the range of day 7 iEAN numbers defined by mean ± SEM of a large set of control C57BL/6J mice. Data is from one representative (A-G and I-O) or 2 pooled independent experiments with 3-5 mice per condition (H). Error bars indicate SD, ns – not significant, *p ≤ 0.05, **p ≤ 0.01, ****p ≤ 0.0001 (unpaired Student’s t test or ANOVA with Tukey’s post-hoc test).

Eosinophils are known to be major producers of IL-4 and IL-13, both of which can skew macrophages towards a tissue-protective phenotype (Liang et al., 2011; Wu et al., 2011). To evaluate the role of these cytokines produced by eosinophils in neuroprotection, we utilized *Epx*^Δ*<u>Il4</u>;Il13*^ mice, in which *Il4* and *Il13* genes are excised by eosinophil peroxidase promoter-driven *Cre* recombinase (Doyle et al., 2013). While *Sv*-induced eosinophilia was preserved in *Epx*^Δ*<u>Il4</u>;Il13*^ mice, subsequent *spiB* infection resulted in significant iEAN loss compared to wild-type control mice (Figures 5G and 5H). Consistently, absence of IL-4 and IL-13 in eosinophils resulted in decreased Arg1 expression by MMs and increased GI transit time (Figures 5I and 5J). To investigate the exact localization of neuroprotective eosinophils, we performed a series of adoptive transfer experiments. Eosinophils derived from the ileum or duodenum from *Sv*-infected mice were transferred to congenic hosts that were subsequently challenged with *spiB* (Figure 5K). Transfer of 6,5×10^5^ duodenum eosinophils resulted in iEAN protection (Figures 5L and 5M). In contrast, equal numbers of ileum eosinophils failed to induce a neuroprotective effect upon spiB challenge; these effects correlated with increased expression of IL-4 by eosinophils derived from duodenum *versus* ileum of *Sv*-infected mice (Figures 5N and 5O). These results suggest that secretion of IL-4 and/or IL-13 by activated duodenum eosinophils mediates distal neuroprotective effects by inducing Arg1 expression in MMs.

Finally, we asked whether long-term neuronal protection is dependent on constant replenishment of eosinophils and if newly recruited macrophages can acquire a tissue-protective phenotype. Following *Sv* infection, we treated mice with anti-CSF1R antibody and waited for new monocytes to be recruited to the tissue and differentiate into MMs (Gabanyi et al., 2016). Despite MM depletion post-*Sv* infection, subsequent *spiB* infection did not result in iEAN loss or reduced Arg1 expression by MMs when compared to isotype-treated animals, suggesting that long-term iEAN protection depends on eosinophils that can induce a protective phenotype in newly developed MMs (Figures S4C-S4E).

Eosinophils were shown to be able to survive up to 3 weeks within tissues, while *Sv*-infection resulted in neuroprotection up to 24 weeks. Based on recent reports (Kaufmann et al., 2018; Mitroulis et al., 2018), we hypothesized that initial *Sv* infection modulates bone marrow hematopoietic stem and progenitor cells, skewing them towards eosinophilic lineage. We analyzed bone marrow of *Sv*-infected mice and found that hematopoietic progenitors (LSK) cells, granulocyte macrophage progenitor (GMP) and eosinophil progenitor (EoP) cells were enriched up to 8 weeks post-infection (Figures S4F-S4J). These results indicate that *Sv* induces long-term restructuring of bone marrow precursors that can sustain an eosinophilic environment, dictating a protective phenotype in tissue macrophages.

## Discussion

Tissues can vary in their susceptibility to damage, regenerative capacity of their cells and in the impact caused by damage to host fitness; consequently, distinct disease tolerance mechanisms have been reported in different tissues and organs (Ayres, 2020; Martins et al., 2019; Medzhitov et al., 2012). The intestine is exposed to high amounts of microbes and potentially harmful substances, and consequently hosts highly regenerative cells like enterocytes, but also cells with low regenerative capacity such as neurons. While an increased regenerative capacity has been reported for enteric neurons, when compared to central nervous system (CNS) neurons (Furness, 2012), infection-induced neuronal loss can have a long-term impact in gut motility and host physiology (Balemans et al., 2017; Matheis et al., 2020; White et al., 2018).

Our findings raise the possibility that infections early in life determine the number of enteric neurons in adult life. Pet store mice had significantly longer small intestines, which has been shown to depend on elevated levels of systemic IL-4 and -13 and in models of chronic helminth infections (Schneider et al., 2018; Wong et al., 2007). Based on intestinal surface area and average neuronal densities, estimated total number of enteric neurons in the ileum is comparable between SPF and pet store mice, indicating that free-living animals might maintain the state of tissue tolerance due to constant exposure to different pathogens. Further studies are needed to define the impact of infections on iEAN during development, as well as the neurochemical code of neurons lost, and restored during episodes of microbe exposure throughout the life of animals (Furness, 2012).

The mechanistic studies described here indicate that the intestinal tissue co-opted pathways induced by distinct pathogens into a convergent tissue macrophage phenotype that mediates enteric neuronal protection, aiding host fitness via intestinal motility (Figure S4). Additionally, initial infections with enteric pathogens used here never affected subsequent bacterial burden and pathogen clearance. Thus, these observations fit the main defining characteristics of disease tolerance (Ayres, 2020; Martins et al., 2019; Medzhitov et al., 2012). The relatively short lifespan of eosinophils does not preclude their involvement in maintaining this state in the long term. This effect was most likely due to epigenetic modulation of myelopoiesis progenitors skewing them towards eosinophilic lineage. Of note, similar mechanisms have until now been mostly described in macrophages and their involvement in disease resistance (Kaufmann et al., 2018; Mitroulis et al., 2018). We hypothesize that certain infections can ‘educate’ HSCs in bone marrow resulting in increased eosinophilic output. As suggested by our comparison between duodenum *versus* ileal eosinophils, the local microenvironment primed by initial infection might then lead to additional eosinophil activation and secretion of cytokines that modulate tissue macrophages. While the impact of disease tolerance mechanisms for host fitness have been demonstrated against a wide range of pathogens, our observations indicate that disease tolerance induced upon infection can have a long-term effect that benefit the host upon subsequent encounter with unrelated pathogens.

## Author Contributions

T.A. initiated, designed, performed the research and wrote the manuscript. B. A., F.M. and C.C. designed and performed experiments. G.F., S.L. generated the cpa3^DTR-tdTomato^ mice. D.M. conceived, initiated, designed, supervised the research, and wrote the manuscript. All authors revised and edited the manuscript and figures.

## Acknowledgements

We thank all Mucida Lab members and Rockefeller University employees for their continuous assistance; A. Rogoz and S. Gonzalez for the maintenance of mice and RU Bio-imaging Research Center for assistance with image acquisition analysis. We thank E. Jacobsen for providing iPHIL and Epx^tm1.1(cre)Jlee^ mouse strains, M. Merad for the anti-CSF1R hybridoma, V. Lenon for anti-Anna-1 antibody, S. Galli and K. Matsushita for the *Sv* larvae, I. Brodsky for *Yp*, J. Lafaille for *H. polygyrus* larvae. We also thank Victora and Lafaille labs for fruitful discussions. This work was supported by NIH Transformative R01DK116646 and R01DK126407, Kenneth Rainin Foundation, Food Allergy FARE/FASI Consortium, NIH R01DK126407 (D.M.). T.A. is a Human Frontier of Science Program postdoctoral fellow. B.A. is a Simons Foundation junior fellow.

## Materials and Methods

### Mice

C57BL/6J (000664), CD45.1 B6 (B6.SJL-Ptprc^a^ Pepc^b^/BoyJ), LysM^Cre^ (B6.129P2-Lyz2^tm1(cre)Ifo^/J, Arg^flox/flox^ (C57BL/6-Arg1^tm1Pmu^/J) and Il4;Il13^flox/flox^ (B6.129P2(Cg)-Il4/Il13^tm1.Lky^/J) were purchased from The Jackson Laboratories and maintained in our facilities. Adrb2^flox/flox^ (Adrb2^tm1Kry^) were generously provided by G. Karsenty, eoCRE (Epx^tm1.1(cre)Jlee^) and iPHIL (Epx^tm2.1(HBEGF)Jlee^) by E. Jacobsen, Hobit-Blimp-1 DKO (Hobit^-/-^Blimp^flox/flox^LCKCre^tg/+^) by K. van Gisbergen. cpa3^DTR-tdTomato^ mice were generated as described below. Pet store mice were purchased at a local pet store (Petco). Mouse lines were interbred in our facilities to obtain the final strains described in the text. Genotyping was performed according to the protocols established for the respective strains by The Jackson Laboratories or personal communication with the donating investigators. Mice were maintained at the Rockefeller University animal facilities under specific pathogen-free conditions. Mice were fed a standard chow diet and used at 7-11 weeks of age for most experiments. Animal care and experimentation were consistent with NIH guidelines and were approved by the Institutional Animal Care and Use Committee at the Rockefeller University.

### Infections

#### *Salmonella enterica* Typhimurium

For infections with Salmonella *spiB*, mice were pre-treated with a single dose of streptomycin (20 mg/mouse dissolved in 100 μl of DPBS) administered by oral gavage 18-24 h prior to infection. Mice were then orally inoculated with 10^9^ CFU of *spiB*. A single aliquot of *spiB* was grown in 3 ml of LB overnight at 37 °C with agitation. Bacteria were then subcultured (1:300) into 3 ml of LB for 3.5 h at 37 °C with agitation and diluted to final concentration in 1 ml of DPBS.

#### Yersinia pseudotuberculosis

*Y. pseudotuberculosis* (strain IP32777) was grown as previously described (Fonseca et al., 2015). Briefly, a single aliquot of Yp was grown in 3 mL of 2xYT media overnight at 28 °C with vigorous agitation and diluted (1:10) to final concentration in DPBS. Mice were fasted for 12-16 h prior to infection with 10^8^ CFU by oral gavage.

#### Strongyloides venezuelensis

*S. venezuelensis* was maintained in our facility in NSG mice by subcutaneous infection with 1000 stage 3 (L3) larvae, resulting in chronic infection of this strain. For each experiment, feces of infected NSG mice were collected and spread on Whatman paper, which was places into a beaker with water and incubated at 28 °C for 2-3 days. Mice were infected subcutaneously with 700 L3 larvae in 200 μl water per mouse. *S. venezuelensis* was passaged periodically by infecting naïve adult NSG mice.

#### Heligmosomoides polygyrus

Mice were infected by oral gavage with 200 third-stage larvae of H. polygyurs in 100 μl water.

### Generation of *cpa3*^DTR-tdTomato^ mice

*Cpa3*^DTR-tdTomato^ knock-in mice were generated using CRISPR/Cas9 technology directly in C57BL/6 mice as described before (REF). Briefly, two guide RNA (gRNA) targeting exon 11 in the 3’ UTR of the cpa3 locus were designed using the online CRISPR Design Tool. These gRNAs were cloned into a plasmid (Addgene) containing Cas9n. The plasmid was then co-injected into the pronucleus with a repair template plasmid (i.e. targeting vector, Addgene) containing an IRES-DTR-tdTomato fusion cassette. WT and knock-in genotypes were confirmed by PCR. DTR and IRES expression were confirmed using the repair template plasmid as a positive control. Insertion of the cassette into the correct locus was verified by using primers located inside the cassette and outside of the 3’ and 5’ homology regions, respectively.

### Antibodies

#### Flow cytometry and whole-mount immunofluorescence antibodies

Antibodies against CD64-APC (Clone X54-5/7.1, cat. # 13906), CD4-BV605 (Clone RM4-5, cat. # 100548), CD150 (Clone TC15-12F12.2, cat. # 115921), CD48-APC (Clone HM480-1, cat. # 103411), Lineage Cocktail-PB (cat. # 133305) were purchases from BioLegend. Antibodies against CD11b-APC-eFluor780 (Clone M1/70, cat. # 47-0112-82), FceR1-APC (Clone MAR1, cat. # 17-5898-80), CD45-PE-Cy7 (Clone 30-F11, cat. # 25-0451-82), CD45-AF700 (Clone 30-F11, cat. # 56-0451-82), CD45.1-PE-Cy7 (Clone A20, cat. # 25-0453-82), GATA-3-PE (Clone TWAJ, cat. # 12-9966-42), FOXP3-eFluor450 (Clone FJK-16s, cat. # 48-5773-82), MHC II-AF700 (Clone M5/114.15.2, cat. # 56-5321-82), Ly-6G-eFluor450 (Clone RB6-8C5, cat. # 48-5931-82), CD11c-AF488 (Clone N418, cat. # 53-0114-82), IL-4-APC (Clone 11B11, cat. # 17-7041-82), Arg1-PE-Cy7 (Clone A1exF5, cat. # 25-36-97-82), Ly6A/E(Sca-1)-PE (Clone D7, cat. # 12-5981-82), CD16/CD32-FITC (clone 93, cat. # 11-0161-82) were purchases from Thermo Fisher Scientific. Antibodies against CD11b-FITC (Clone M1/70, cat. # 553310), Siglec-F-APC-Cy7 (Clone E50-2440, cat. # 565527), -PE (Clone E50-2440, cat. # 552126) and -BV421 (Clone E50-2440, cat. # 56268), c-kit-PE-Cy7 (Clone 2B8, cat. # 558163), CD45.2-PerCP-Cy5.5 (Clone 104, cat. # 552950), CD45.2-AF700 (Clone 104, cat. # 560693), CD45R-FITC (Clone RA3-6B2, cat. # 553088), CD8a-AF488 (Clone 53-6.7, cat. # 557668), I-A/I-E-FITC (Clone M5/114.15.2, cat. # 553623), CD125-AF488 (Clone T21, cat. # 558533) were purchased from BD. Cell surface and intracellular antibodies were used at 1:200 and 1:100 dilution, respectively. Antibody against ANNA-1 was a gift from Dr. Vanda A. Lennon. Fluorophore-conjugated secondary antibody H&L goat anti-human-AF568 (Thermo Fisher Scientific) was used to detect anti-ANNA-1 antibody.

#### Antibodies for in vivo experiments

Antibodies against CD4 (Clone GK1.5, cat. # BE0003-1), CSF1R (Clone ASF98, cat. # BE0213) and IL-5 (Clone TRFK5, cat. # BE0198) and corresponding isotype controls were purchased from BioXCell. CD4 mAb was injected i.p. on indicated days at 200 μg/mouse/timepoint. IL-5 mAb was injected i.p. on indicated days at 500 μg/mouse/timepoint. CSF1R mAb was injected i.p. on indicated days at 50 mg/kg. Mice were previously habituated to i.p. injections for at least 5 days prior to the antibody treatments.

### Ivermectin treatment

Ivermectin (Milipore Sigma) was dissolved in DMSO at 100 mg/ml. 14 days after *Sv* infection, ivermectin was orally gavaged at 50 μg/mouse. 14 days after *Hp* infection ivermectin was injected subcutaneously at 20 mg/kg.

### Diphtheria toxin treatment

DT (Sigma Aldrich) was reconstituted in DPBS. iPHIL mice were treated i.p. on indicated days at 15 ng/gram body weight. Cpa3-DTR mice were treated i.p. on indicated days at 300 μg/mouse.

### Intestine dissection

Mice were sacrificed and duodenum (1 cm moving distal from the gastroduodenal junction) or ileum (1 cm moving proximal from the ileocecal junction) was removed. For dissection of the *muscularis*, following the above procedures, the intestinal tissue was placed on a chilled aluminum block with the serosa facing up (Gabanyi et al., 2016). Curved forceps were then used to carefully remove the *muscularis*. Tissue was then used for whole-mount imaging or flow cytometry.

### *Muscularis* processing for flow cytometry

*Muscularis* was finely cut and digested in HBSS Mg^2+^Ca^2+^ + 5% FBS + 1x NaPyr + 25mM HEPES + 50 μg/ml DNaseI (Roche) + 400 U/ml Collagenase D (Roche) + 2.5 U/ml Dispase (Corning) at 37 °C. The *muscularis* was digested for 40 min. The tissue was then homogenized with an 18-gauge needle and filtered through a 70 μm cell strainer and washed with HBSS Mg^2+^Ca^2+^. The cells were incubated with Fc block and antibodies against the indicated cell surface markers in FACS buffer (PBS, 1% BSA, 10 mM EDTA, 0.02% sodium azide).

### Bone marrow processing

Bone marrow cells from femurs were passed through 70 μm filter and treated with red blood cell lysis buffer, followed by staining with fluorescently labeled antibodies.

### Isolation of intraepithelial and lamina propria cells

Intraepithelial and lamina propria cells were isolated as previously described (Bilate et al., 2016). Briefly, sections of small intestines were harvested and washed in PBS and 1mM dithiothreitol (DTT) followed by 30 mM EDTA. Intraepithelial cells were recovered from the supernatant of DTT and EDTA washes and mononuclear cells were isolated by gradient centrifugation using Percoll. Cells from lamina propria were obtained after collagenase 8 digestion of the tissue. Single-cell suspensions were then stained with fluorescently labeled antibodies for 30 min on ice.

### Intracelluar staining for flow cytometry

Intranuclear staining for transcription factors was conducted using Foxp3 / Transcription Factor Staining Buffer Set according to manufacturer’s instructions (eBioscience, USA). Intracellular staining for cytokines was conducted in Perm/Wash buffer after fixation and permeabilization in Fix/Perm buffer (BD Biosciences, USA) according to kit instructions. Flow cytometry data were acquired on an LSR-II flow cytometer (Becton Dickinson, USA) and analyzed using FlowJo software package (Tri-Star, USA).

### Gating strategies

For flow cytometric analysis following gating strategy was used to identify macrophages: single, live, myeloid cells (based on FSC, SSC and live/dead fixable dye Aqua stain), CD45+, CD11b+ and CD64+. Eosinophils: single, live, myeloid cells, CD45+, CD11b+ and Siglec-F+. Mast cells: single, live, myeloid cells, CD45+, CD117+ and FceR1+. For all cell types, B220+ and CD8a+ cells were excluded.

### Eosinophil sorting

Following lamina propria cell isolation, cells were processed as previously described (Geslewitz et al., 2018). Cells were then sorted using a FACSAria™ cell sorter flow cytometer (Becton Dickinson) and subsequently transferred by retro-orbital injection at numbers indicated in Figure 5.

### Cytokine quantification

After centrifugation of peripheral blood, the serum samples were immediately aliquoted and stored at 4 °C until the experiment was conducted. The concentrations of IL-4 and IL-13 were measured once after collection of each individual samples by using the cytometric bead array assay (BD Biosciences) according to the manufacturer’s instructions.

### CFU counting

Fecal pellets from *spiB- or Yp-*infected mice were weighed and then disrupted in 400 mL of DPBS. Serial dilutions were made from the original suspension and then 5 ml of each dilution was plated onto Salmonella-Shigella or 2xYT-Irgasan plates, respectively. The plates were then incubated overnight, and the number of black colonies were counted for the serial dilution with the clearest delineation of single units. This number was then multiplied by the dilution factor and by 80 to give the number of colony-forming units (CFU) in the original suspension. CFU numbers were then divided by the original fecal pellet weight to give the number of CFU per mg of feces.

### Quantification of *Sv* eggs in feces

Fecal pellets from *Sv-*infected mice were weighed and then disrupted in 400 μl of DPBS. 100 μl of iodine solution was added to increase the visibility of fecal eggs. Eggs in small portions of each sample were counted under a microscope, and the number of eggs per gram of feces was determined for each sample.

### Intestine motility measurements

For measurement of total intestinal transit time, mice were given an oral gavage of 6% carmine red (Sigma-Aldrich) dissolved in 0.5 % methylcellulose (prepared in sterile 0.9% NaCl). Total intestinal transit time was measured as the time from oral gavage it took for mice to pass a fecal pellet that contained carmine.

### Neuronal imaging and counting quantification

After dissection muscularis was pinned down on a plate coated with Sylgard and then fixed overnight with 4 % PFA. After washing in DPBS whole mount samples were then permeabilized first in 0.5 % Triton X-100 (perm buffer) for 2 h at room temperature (RT) with gentle shaking. Samples were then blocked for 2 h in blocking buffer (5 % bovine serum albumin and 5 % goat serum in perm buffer) for 2 h at RT with gentle agitation. Anti-ANNA-1 antibody was added to the blocking buffer at appropriate concentrations and incubated for 2 days at 4°C. After primary incubation the tissue was washed 4 times in perm buffer and then incubated in blocking buffer with secondary antibody for 2-3 h at RT. Samples were again washed 4 times in perm buffer and then mounted with FluoroMount G on slides with coverslips. Slides were kept in the dark at 4 °C until they were imaged. A minimum of 5-10 images were randomly acquired across a piece of whole-mount *muscularis*. These images were then opened in Fiji, and the cell counter was used to count the number of ANNA-1+ cells in a given field. This number was then multiplied by a factor of 2.95 (20x objective) or 3.125 (25x objective), to calculate the number of counted neurons per square millimeter (mm^2^). The average of 8-10 images was then calculated and plotted. Wholemount intestine samples were imaged on an inverted LSM 880 NLO laser scanning confocal and multiphoton microscope (Zeiss) and on an inverted TCS SP8 laser scanning confocal microscope (Leica).

### CG-SMG dissections and cFos staining

Mice were cervically dislocated and a midline incision was made; the viscera were removed out of the peritoneal cavity. The superior mesenteric artery was located by identifying the intersection of the descending aorta and left renal artery. Fine forceps and microdissection scissors were used to remove the CG-SMG which is wrapped around the superior mesenteric artery and associated lymphatic vessels. CG-SMG samples were washed in cold 1x PBS and fixed overnight at 4 °C (rotating) in fresh 4% PFA. CG-SMG samples were then washed four times in 1x PBS at room temperature and blocked overnight in 5% Normal Donkey Serum with 0.5% Triton X-100/0.05% Tween-20/4 μg heparin (PTxwH) at 4 °C. Samples then incubated with anti-cFos (1:500; Cell Signaling Technologies, 2250S) and anti-TH (1:200; Milipore; AB1542) for 48 hours at 4 °C (rotating). Samples were washed four times in PTxwH at room temperature and incubated with highly cross-adsorbed donkey anti-rabbit AF647 (Invitrogen A31573) and donkey anti-sheep AF568 (Invitrogen; A21099) antibodies at 4 °C for 48 hours (rotating). Samples were washed four times in PTxwH at room temperature and mounted in Fluoromount G. TH staining was used to identify the sympathetic neurons and number of cFos+ nuclei was counted in multiple z-stack images with confocal microscope. ImageJ Cell Counter plugin was used to count the cFos+ nuclei and data were not normalized to area or volume. Each data point represents the total number of cFos+ cells per CG-SMG.

### Intestine neuronal quantification: neurons per ganglion

A myenteric ganglion was defined as a continuous group of ANNA1+ cells that are separated by less than 15 μm in distance. Only complete ganglia were counted per field of view. Thus, the following ganglia were excluded: 1. Ganglia that were truncated; 2. No clear separation (>15 μm) was noted between the last ANNA1+ cell and the edge of the field of view. In the case of single ANNA1+ cells that are separated by 15μm on all sides, this was counted as extraganglionic. The number of quantifiable ganglia was averaged across a minimum of 10 images per gut segment per animal.

### Statistical analysis

Significance levels indicated are as follows: * p < 0.05, ** p < 0.01, *** p < 0.001, **** p < 0.0001. All data are presented as mean ± SD or mean ± SEM. At least two independent experiments were performed throughout in this study. All statistical tests used were two-tailed. Multivariate data were analyzed by one-way ANOVA and Tukey’s multiple comparisons post hoc test. Comparisons between two conditions were analyzed by unpaired Student’s t test. GraphPad PRISM version 9 was used for generation of graphs and statistics.

---

**Figure S1.**
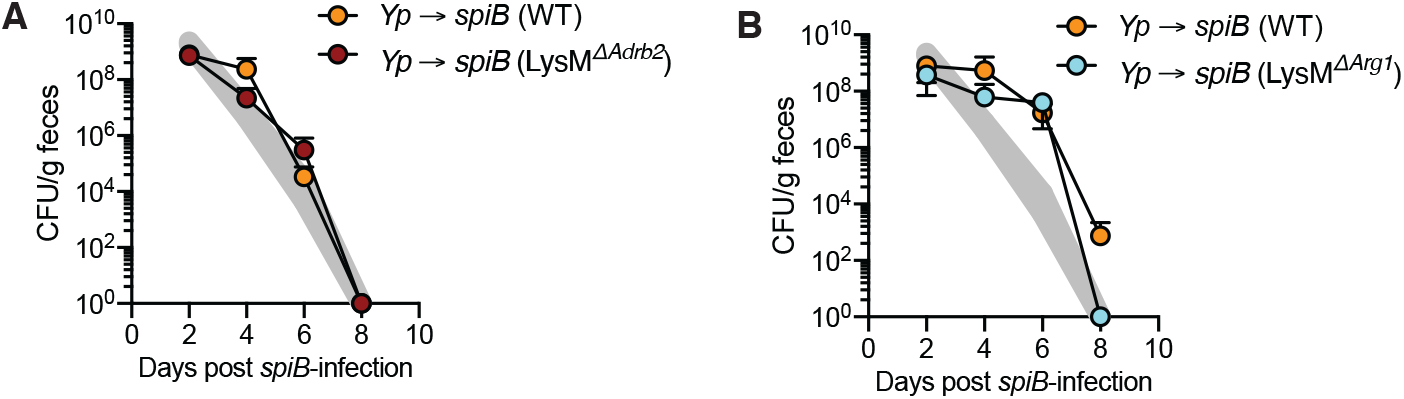
Initial *Yp* infection does not impact subsequent *spiB* clearance in LysM^*ΔAdrb2*^ or LysM^*ΔArg1*^ mice. (A and B) LysM^*ΔAdrb2*^, LysM^*ΔArg1*^ mice and WT littermates were orally gavaged with *Y. pseudotuberculosis* and 21 days later with *Salmonella typhimurium spiB*. Quantification of fecal CFU in LysM^ΔAdrb2^ (A) and LysM^ΔArg1^ (B) mice. Shaded area indicates range of *spiB* CFU numbers defined by mean ± SEM of a large set of control C57BL/6J mice.

**Figure S2.**
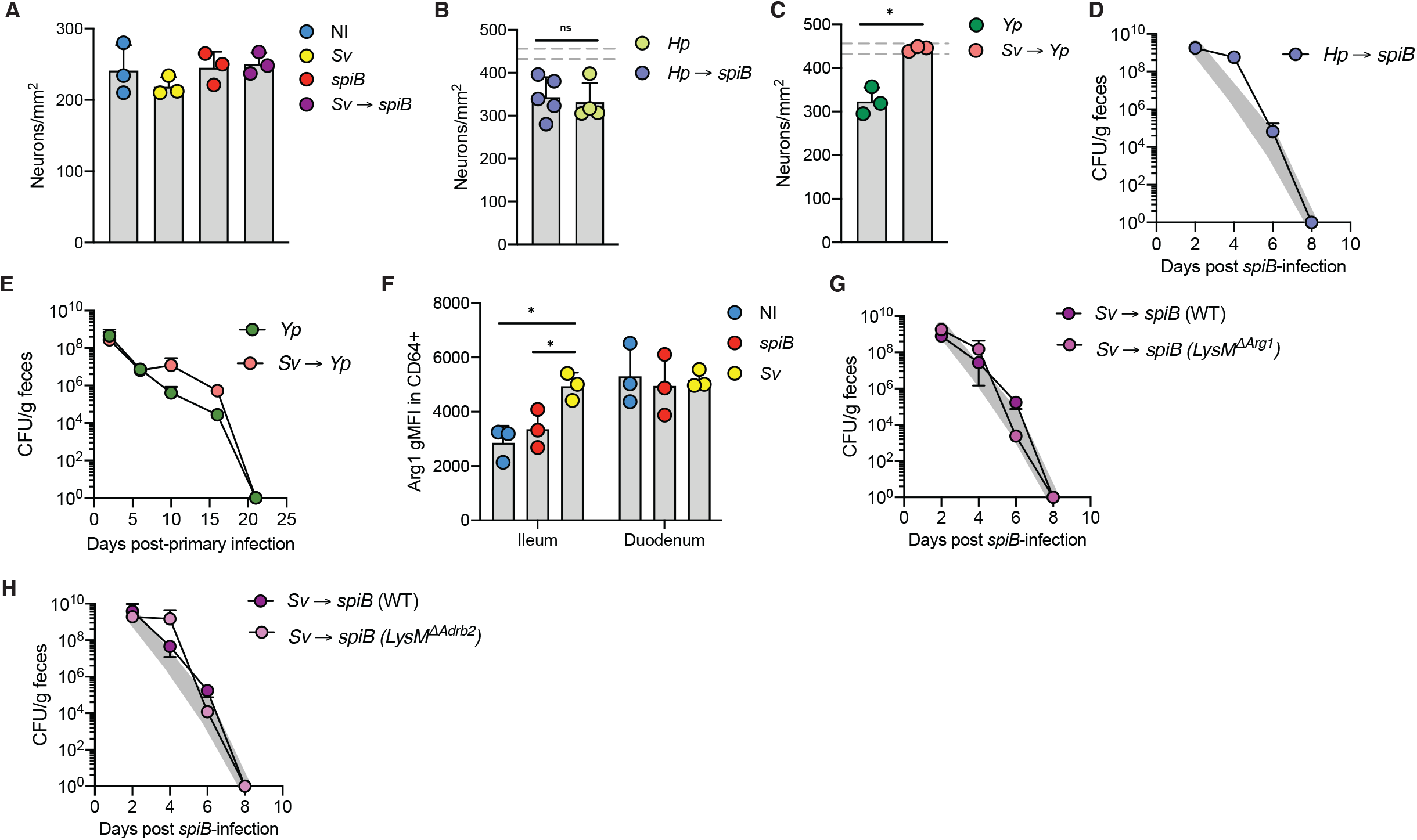
Helminth infections induce tissue tolerance. (A) C57BL/6J mice were s.c. injected with water and then orally gavaged with PBS (non-infected, NI), 10^9^ colony-forming units (CFU) of *Salmonella spiB*, or subcutaneously injected with 700 *S. venezuelensis* (Sv) larvae only or 700 *Sv* larvae and 14 days later with 10^9^ CFU of *spiB*. (A) Neuronal quantification in the duodenum myenteric plexus day 8 post-*spiB* infection. (B and C) C57BL/6J mice were orally gavaged with 200 *H. polygyrus* (*Hp*) larvae only or *Hp* larvae and 14 days later with *spiB*. (B) Quantification of ileum myenteric plexus neurons at 7 days post-*spiB* infection. (C) Quantification of fecal CFU. (D and E) C57BL/6J mice were orally gavaged with *Yp* or subcutaneously injected with *Sv* followed by *Yp* 14 days later. (D) Quantification of ileum myenteric plexus neurons at 21 days post-*Yp* infection. (E) Quantification of fecal CFU. (F) Frequencies of Arg1-expressing ileum and duodenum myenteric plexus macrophages in C57BL/6J mice orally gavaged with PBS (non-infected, NI), 10^9^ colony-forming units (CFU) of *Salmonella spiB*, or subcutaneously injected with 700 *S. venezuelensis* (*Sv*) larvae. Flow cytometry analysis was performed at 14 days-post infection. (G and H) LysM^Δ*Adrb2*^, LysM^Δ*Arg1*^ mice and WT littermates were infected with *Sv* and 14 days later with *spiB*. Quantification of fecal CFU in LysM^Δ*Arg1*^ (G) and LysM^Δ*Adrb2*^ (H) mice. (C, G and H) Shaded area indicates range of *spiB* CFU numbers defined by mean ± SEM of all control C57BL/6J mice. (B and D) Dashed lines indicate the range of day 7 iEAN numbers defined by mean ± SEM of a large set of control C57BL/6J mice. All data is representative of 2 independent experiments with 3-5 mice per condition. Error bars indicate SD, ns – not significant, * p ≤ 0.05 (unpaired Student’s t test or ANOVA with Tukey’s post-hoc test).

**Figure S3.**
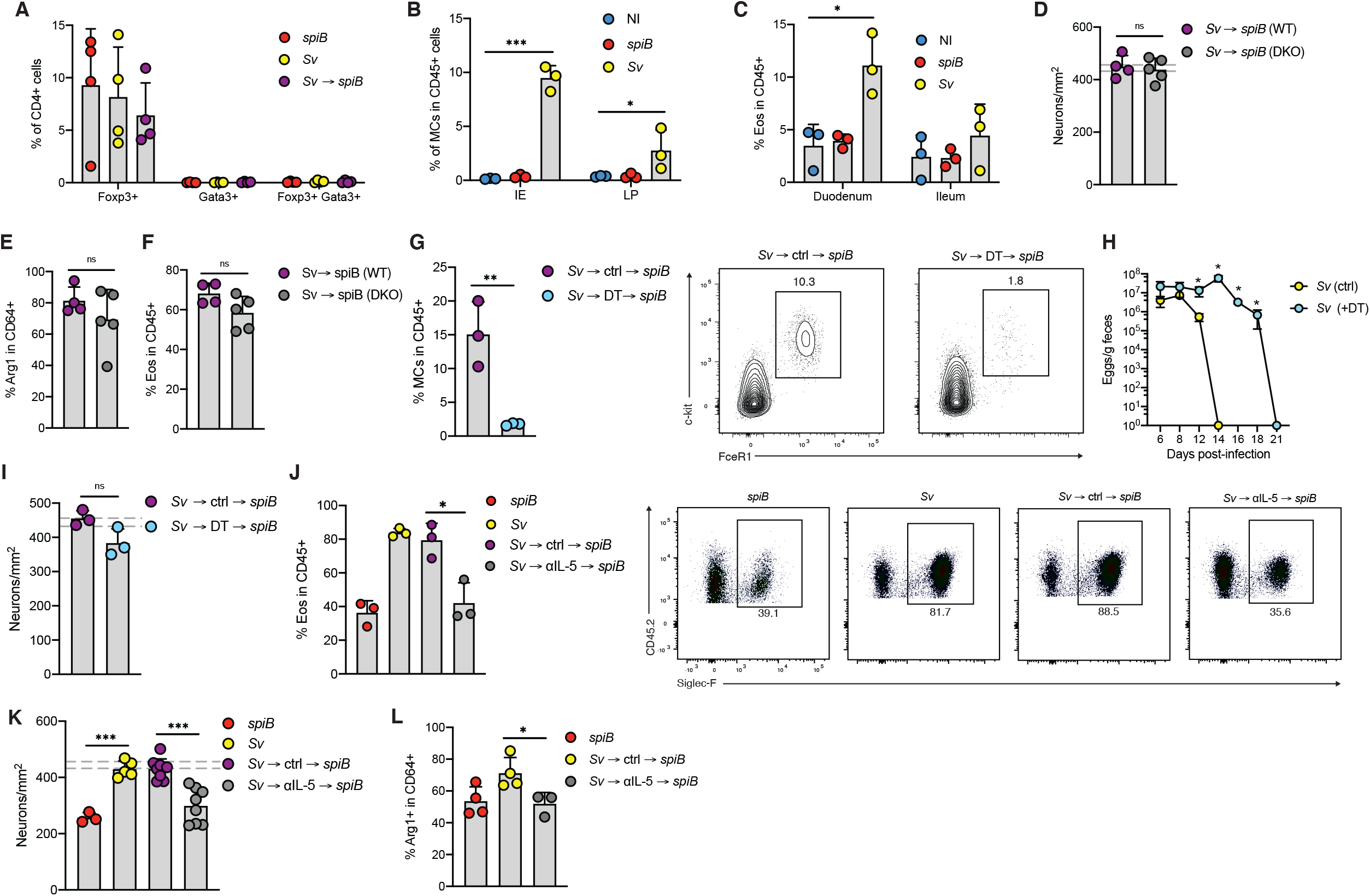
Helminth induced neuroprotection depends on IL-5. (A-C) C57BL/6J mice were s.c. injected with water (non-infected, NI) or infected with *spiB, Sv* only or *Sv* and 14 days later with *spiB*. (A) Flow cytometry intranuclear analysis of CD4^+^ T cells in ileum lamina propria (LP) expressing indicated transcription factors 2 days post-*spiB* infection. (B) Frequencies of duodenum IE and LP mast cells at 14 dpi with *Sv*. (C) Frequencies of duodenum and ileum IE eosinophils at 14 dpi with *Sv*. (D-F) Hobit and Blimp-1 DKO (*Hobit^-/-^Blimp1^flox/flox^LckCre^tg/+^*) mice and WT littermates were infected with *Sv* and 28 days later with *spiB*. (D) Quantification of ileum myenteric plexus neurons at 7 dpi with *spiB*. (E) Arg1 expression by ileum myenteric plexus macrophages at 7 dpi. (F) Frequency of duodenum LP eosinophils at 7 dpi. (G-I) cpa3^DTR-tdTomato^ mice and WT littermates were infected with *Sv* and treated with DT on day 0, 1, 4, 7, 10, 13, 16 and 18 post-infection. On day 21 post-*Sv* infection mice were orally gavaged with *spiB*. (G) Frequency of duodenum IE mast cells at 7 dpi with *spiB*. (H) Quantification of fecal *Sv* eggs. (I) Quantification of ileum myenteric plexus neurons at 7 dpi. (J-L) C57BL/6J mice were infected with *spiB* or *Sv*. *Sv*-infected mice were treated with anti-IL-5 antibody on day 0, 4, 8, 11 and 14 followed by oral gavage with *spiB*. Data is from one representative (A-D) or 2 pooled independent experiments with 3-5 mice per condition (G-J). Error bars indicate SD, ns – not significant, * p ≤ 0.05 (unpaired Student’s t test or ANOVA with Tukey’s post-hoc test).

**Figure S4.**
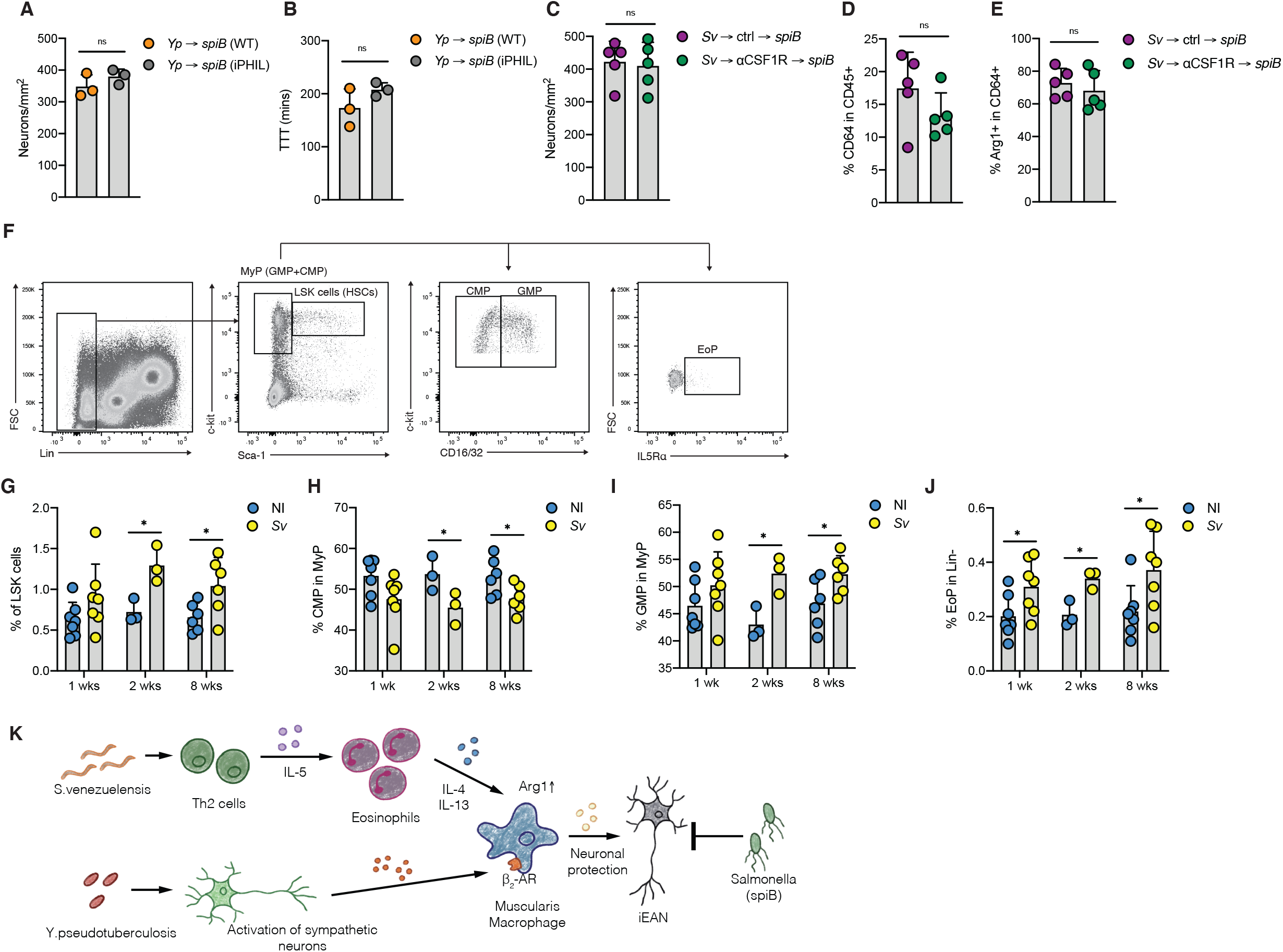
*S. venezeluensis* infection induces bone marrow hematopoietic progenitors. (A and B) iPHIL mice and WT littermates were infected with *Yp* and treated with DT on day 0, 1, 4, 7, 10, 13, 16 and 18 post-infection. On day 21 post-*Sv* infection mice were orally gavaged with *spiB*. (A) Quantification of ileum myenteric plexus neurons at 7 dpi. (B) Total GI transit time measured 10 days *post-spiB* infection. (C-E) C57BL/6J mice were infected with *Sv* and at 14 dpi treated with anti-CSF1R or IgG isotype control antibodies, followed by oral gavage with *spiB* 14 days later. (C) Ileum myenteric plexus neurons were quantified at 7 days post *spiB*-infection. (D) Frequency of ileum myenteric plexus macrophages at 7 dpi. (E) Frequency of Arg1-expressing ileum myenteric plexus macrophages at 7 dpi with *spiB*. (F-J) Frequencies of indicated bone marrow progenitor cell subsets 1, 2 and 8 weeks post-*Sv* infection. (K) Graphical representation of main findings. Bacteria-induced neuroprotection relied on β_2_-adrenergic receptor signaling in MMs. Helminth-induced neuroprotection was dependent on T cells and systemic production of interleukin (IL)-4 and -13 by eosinophils, which regulate MMs located in a different intestinal region. These two pathways converged into Arginase-1 positive MMs that provide neuroprotection against subsequent infection.Data is from one representative (A-E) or 2 pooled independent experiments with 3-5 mice per condition (G-J). Error bars indicate SD, ns – not significant, * p ≤ 0.05, **** p ≤ 0.0001 (unpaired Student’s t test or ANOVA with Tukey’s post-hoc test).

